# Improved fluorescent proteins for dual-color post-embedding CLEM

**DOI:** 10.1101/2022.02.16.480633

**Authors:** Dingming Peng, Na Li, Wenting He, Kim Ryun Drasbek, Tao Xu, Mingshu Zhang, Pingyong Xu

**Affiliations:** Key Laboratory of RNA Biology, Institute of Biophysics, Chinese Academy of Sciences, Beijing, China; Sino-Danish College, University of Chinese Academy of Sciences, Beijing, China; Sino-Danish Center for Education and Research, Beijing, China; Bioland Laboratory (Guangzhou Regenerative Medicine and Health Guangdong Laboratory), 510005, Guangzhou, China; Center of Functionally Integrative Neuroscience, Dept of Clinical Medicine, Aarhus University, Aarhus, Denmark; National Laboratory of Biomacromolecules, CAS Center for Excellence in Biomacromolecules, Institute of Biophysics, Chinese Academy of Sciences, Beijing, China; College of Life Sciences, University of Chinese Academy of Sciences, Beijing, China

**Keywords:** Dual-color CLEM, high SBR, RSFP, probe development

## Abstract

Post-embedding correlative light and electron microscopy (CLEM) has the advantage of high-precision registration and enables light and electron microscopy imaging of the same slice. However, its broad application has been hampered by the limited available fluorescent proteins (FPs) and low signal-to-background ratio (SBR). Here, we developed a green photoswitchable FP, mEosEM-E with substantially high on/off contrast in EM samples embedded in Epon resin which maximally preserves cellular structures but quenches the fluorescence of FPs. Taking advantage of the photoswitching property of mEosEM-E, the autofluorescence background from the resin was significantly reduced by a subtraction-based CLEM (sCLEM) method. Meanwhile, we identified a red fluorescent protein (RFP) mScharlet-H that exhibited higher brightness and SBR in resin than previously reported RFPs. With mEosEM-E and mScharlet-H, dual-color post-Epon-embedding CLEM images with high SBR and no cross-talk signal were successfully performed to reveal the organization of nucleolar proteins. Moreover, a dissection of the influences of different EM sample preparation steps on the fluorescence preservation for several RFPs provides useful guidance for further probe development.

## Introduction

Light microscopy (LM) highlights and discriminates one or a few biomolecules using fluorescent labeling. Electron microscopy (EM), on the other hand, reveals the ultrastructural context of the cell where the biomolecules reside. Correlative light and electron microscopy (CLEM) [1] integrates the information of both LM and EM in a complementary way, and thus provides localization, structural and functional insights into the biomolecular machines, and shows great application prospects in cellular physiology [2], virology [3] and neuroscience [4]. In CLEM imaging, LM can not only indicate the location of the target protein, but also be used as a guide to quickly find slices containing target cells or proteins in a large number of EM slices, and to quickly locate the cell on the target slice for EM imaging. The advantage of fluorescence navigation is unique and is not possessed by EM labeling techniques, such as immunogold labeling, APEX-gold [5], APEX2 [6], miniSOG [7], and other metal [8] or chemical [9] tags.

Same as any other imaging technique, the practical performance of CLEM is largely determined by the labeling probe. However, most fluorescent proteins (FPs) experienced severe fluorescence loss during standard EM sample preparation, making it difficult to perform LM and EM on the same section that can be correlated with high precision. To circumvent this problem, several strategies have been used. Pre-embedding CLEM [10, 11] obtains fluorescence microscopy (FM) image ahead of EM sample preparation, avoiding fluorescence quenching caused by a series of chemical treatments. However, because of the morphology distortion during EM sample preparation and sectioning, it suffers from poor registration between FM and EM images. Several modified EM sample preparation protocols with no or reduced OsO_4_ concentration have been reported to preserve sufficient fluorescence signal for post-embedding CLEM [12, 13], however, deteriorated EM images were often observed.

In 2015, Paez-Segala et al. engineered the first FP, mEos4b, to retain fluorescence after 1% OsO_4_ treatment [14]. However, the hydrophilic GMA resin was used for mEos4b-labeled samples. Compared to the GMA resins, the hydrophobic Epon resin has advantages in maintaining cellular ultrastructure as well as sectioning quality [3, 15] due to its higher toughness and hardness. Nevertheless, Epon resin undermines fluorescence more severely. To solve this problem, we have previously developed mEosEM [16], an FP that survives 1% OsO_4_ fixation and Epon embedding and enables super-resolution CLEM (SR-CLEM) due to the preserved photomodulable property. Recently, Tanida et al. found that mKate2 [17], mCherry2, mWasabi, and GoGFP-v0 [18] could also preserve fluorescence after standard EM sample preparation and dual-color CLEM imaging was achieved using mWasabi and mcherry2. However, both the green and the red channel images showed very low signal-to-background ratio (SBR), making it difficult to distinguish the real fluorescence signals from that of the background or noises.

Low SBR can be attributed to three aspects. First, most of the fluorescence signals of the FPs mentioned above are quenched after TEM sample preparation, and only a small amount of the remaining fluorescence signal is used for CLEM imaging. Second, the thickness of ultrathin sections is generally only about 100 nm, and the number of FP-labeled molecules in the sections is limited. Third, the Epon resin has strong autofluorescence, especially in the green channel (Figure S1). Therefore, developing FPs with high in-resin SBR and repressing the autofluorescence background are effective ways to solve the problem. For dual-color CLEM, another phenomenon worth noting is that red fluorescent proteins (RFPs) on Epon-embedded slices emitted green fluorescence when illuminated with 488-nm laser (Figure S8), which interfered the signal in the green channel. Therefore, FPs and imaging methods that can solve these problems are desirable.

Reversibly photoswitchable FP (RSFP) can be utilized to suppress the unmodulatable fluorescent background and enhance the signal contrast by means of optical lock-in detection (OLID) [19], synchronously amplified fluorescence image recovery (SAFIRe) [20, 21] and out-of-phase imaging after optical modulation (OPIOM) [22]. We speculated that similar strategies could be applied with an OsO_4_ - and Epon-resistant RSFP to eliminate the resin background and the RFP crosstalk signal. In this study, we developed a fluorescence background-reduced CLEM method (sCLEM) using a simple subtraction of images of RSFPs at fluorescent on- and off-states and efficiently extracted the fluorescence signals of the FPs from that of the background. The higher on/off contrast of the RSFP, the better SBR of the final image. Therefore, we evolved a mEosEM variant termed mEosEM-E, with high on-state brightness and on/off contrast ratio after standard EM sample preparation, and demonstrated its utility in sCLEM imaging.

On the other hand, despite a few FPs were reported for post-Epon-embedding CLEM [14, 16–18], all these probes were discovered by OsO_4_ resistance assay, the results of which are sometimes inconsistent with the final performance of the probe in CLEM imaging. As a matter of fact, nearly every step during EM sample preparation will quench the fluorescence of FPs. However, a comparative study that dissects the influence of each step on fluorescence signal reduction is lacking. In order to develop an optimal RFP that can be coupled with mEosEM-E for dual-color CLEM imaging, we assessed the fluorescence preservation of nine commonly used RFPs after each key step of EM sample preparation, including pre-fixation, OsO_4_ fixation, ethanol dehydration, and Epon embedding. The results showed that the OsO_4_ resistance assay is not the optimal criterion for CLEM probe development, while the performance of a probe should be evaluated in the final sample section. mScarlet-H, which preserved the highest on-section fluorescence and SBR among the nine RFPs tested, is the best RFP reported to date for the same section post-Epon-embedding CLEM. Finally, high SBR and accurate dual-color CLEM imaging of nucleolar proteins was successfully achieved for the first time using mEosEM-E and mScarlet-H double labeling.

## Results

### Green mEosEM-E with high SBR for sCLEM

To obtain a CLEM probe with high on/off contrast, we chose mEosEM as the template. mEosEM is a photoconvertible FP (PCFP) that can be converted to an RFP upon 405-nm illumination, but it also has photoswitching property at the green state like an RSFP. After the TEM sample preparation, mEosEM lost the characteristic of light conversion while retaining the photoswitching property[16]. However, this photoswitching property was not used for background removal in CLEM imaging, and the on/off contrast was not optimized for this use.

Our previous study revealed that the first amino acid of the chromophore tripeptide (XYG) of the Eos FP family not only determines the photomodulable type (RSFP or PCFP) of the FP but also greatly affect its photoswitching property [23]. We reasoned that because the first amino acid of the chromophore is located inside the barrel structure of the FP, mutagenesis at this site would have no negative effect on the resistance to the EM sample preparation, while largely tuning the photoswitching contrast. As expected, saturation mutagenesis at His63 of mEosEM produced a series of RSFP mutants displaying a variety of switching contrast and kinetics (Figure S2). We found that among several improved mutants, mEosEM-E (mEosEM H63E) displayed substantially improved contrast, in other words, largely reduced normalized residual fluorescence as compared to mEosEM after TEM sample preparation (Figure 1A). The on/off contrast ratio of mEosEM-E is ∼2.7-fold higher than that of mEosEM on average (Figure 1B).

**Figure 1.**
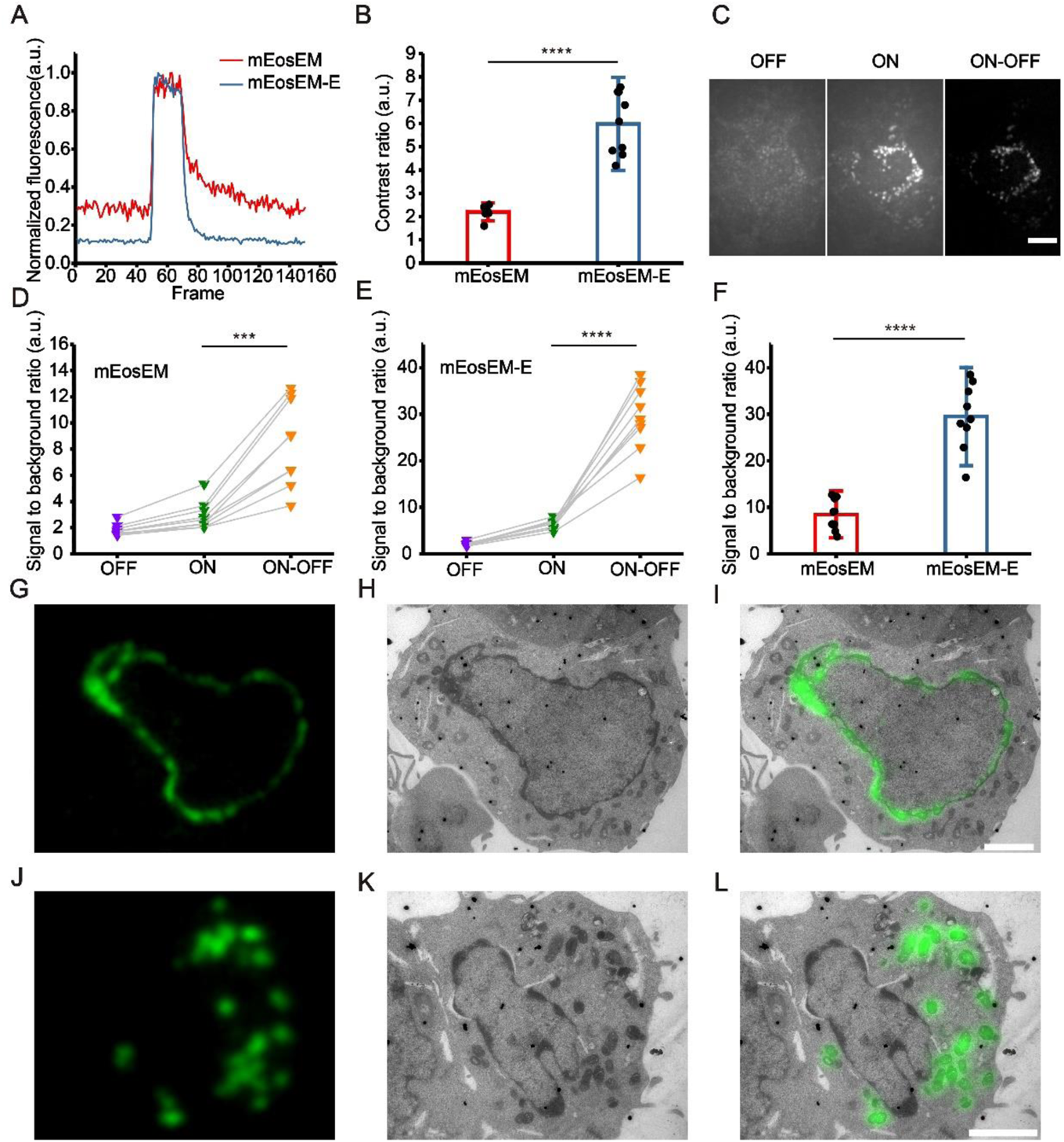
mEosEM-E is an improved green RSFP for high SBR sCLEM imaging. (**A**) Representative normalized photoswitching curves of Epon-embedded HEK 293T cell sections (100 nm) expressing mitochondria-targeted mEosEM (red) and mEosEM-E (blue). Samples were excited and switched off with the 488-nm laser (0.41 kW/cm_2_), and switched on with the 405-nm (0.21 kW/cm_2_) laser for 1 s. (**B**) Statistics photoswitching contrast ratio of mEosEM (red) and mEosEM-E (blue). Data are summarized in Table S1. Bars represent mean ± SD. P values were determined with two-tailed t-test (n = 9). **** indicates p < 0.0001. (**C**) Representative OFF (left), ON (middle), and ON ─ OFF (right) images of Epon-embedded HEK 293T cell sections (100 nm) expressing mitochondria targeting mEosEM-E. Scale bar, 5 µm. (**D, E**) SBR of mEosEM- (**D**) and mEosEM-E- (**E**) labeled cell samples in OFF (purple), ON (green), and ON ─ OFF (orange) images. Data are summarized in Table S2 & 3. P values were determined with two-tailed t-tests (n = 9). *** indicates p < 0.001, **** indicates p < 0.0001. (**F**) Statistics SBR of mEosEM (red) and mEosEM-E (blue) in ON ─ OFF images. Two-tailed t-tests were performed (n = 9). **** indicates p < 0.0001. Data are summarized in Table S4. (**G-L**) Post-Epon-embedding CLEM of mEosEM-E. FM, EM, and CLEM images of nuclear envelope (**G-I**) and mitochondria (**J-L**) labeled by mEosEM-E (100 nm slices). Scale bars, 2 µm.

To demonstrate the superiority of mEosEM-E in CLEM, we imaged HEK 293T cell sections expressing mitochondria-targeting mEosEM-E or mEosEM. Fluorescence image sequences were acquired under the illumination of the 488-nm laser alone followed by the 488- and 405-nm lasers together. We termed the images acquired without the illumination of 405-nm laser as “OFF images” and those with 405-nm laser as “ON images”. Every 20 frames of the OFF and subsequent ON images from the image stack were averaged respectively. Then, the subtraction was performed pixel by pixel between the averaged ON and OFF images to acquire the final “ON ─ OFF image” (Figure S3). We named the subtraction–based LM as sLM, and accordingly CLEM as sCLEM. It can be clearly seen that the autofluorescent background in the OFF image was almost completely removed in the ON ─ OFF image of mEosEM-E-labeled cells (Figure 1C). For both mEosEM and mEosEM-E, the SBR in the ON ─ OFF image was significantly increased compared to the ON image (Figure 1D & E). Notably, the SBR of mEosEM-E fluorescence is 3.46-fold higher than that of mEosEM in the ON ─ OFF image (Figure 1F). Further exploration revealed that the substantial higher SBR in the ON ─ OFF image of mEosEM-E is mainly attributed to its higher SBR in the ON image (Figure S4).

Next, same section post-Epon-embedding sCLEM was demonstrated in HEK 293T cells, of which the nuclear lamina (Figure 1G-I) and the mitochondria matrix (Figure 1J-L) were labeled by mEosEM-E, separately. The results showed that high SBR and nearly background-free fluorescence images were obtained and well aligned with the EM images.

Altogether, mEosEM-E proved to be a better CLEM probe for high SBR imaging and is potentially useful for investigating proteins of low expression level.

### Identification of mScarlet-H as a high-performance red CLEM probe

To find the optimal red probe for dual-color CLEM, nine commonly used RFPs were chosen for investigation: mScarlet, mScarlet-H, mScarlet-I, FusionRed-MQV, mRuby3, mApple, tdTomato, mKate2 [17], mCherry2 [18], of which the latter two were previously demonstrated feasible for post-Epon-embedding CLEM. Instead of performing only the OsO_4_-resistant assay reported in the previous papers [16–18], we thoroughly analyzed the fluorescence preservation after each key step of the EM sample preparation procedure, including aldehyde fixation, OsO_4_ fixation, ethanol dehydration, Epon embedding, and high-temperature polymerization.

A direct comparison of the absolute fluorescence intensity in fixed cells, which is a representative of the practical performance of the probe, showed that pre-fixation with 2% formaldehyde and 2.5% glutaraldehyde already significantly reduced fluorescence intensities of several RFPs, up to 50% of their initial value. While a few others, such as mScarlet-H and FusionRed-MQV were much less affected by pre-fixation, indicating a variated response among different RFPs (Figure S5). The fluorescence intensities of all RFPs were markedly reduced after 1% OsO_4_ fixation, nevertheless, mScarlet-I, mRuby3, mScarlet, and mScarlet-H are the top 4 FPs retaining high residue fluorescence, which are all substantially higher than that of the previously reported mKate2 and mCherry2 (Figure 2A). However, the subsequent dehydration step using 100% ethyl alcohol changed the ordering of RFPs, making only mRuby3 standout, while others diminished to a similar level (Figure 2B). We speculated that the advantage of mRuby3 at this stage is due to the insensitivity of mRuby3 to the dehydration treatment, which was later confirmed by the results from dehydration treatment without OsO_4_ fixation (Figure S6). Notably, the robustness of mRuby3 to the dehydration treatment following 1% OsO_4_ fixation is not as strong as that to a single dehydration treatment, indicating a superimposed effect of the two treatments.

**Figure 2.**
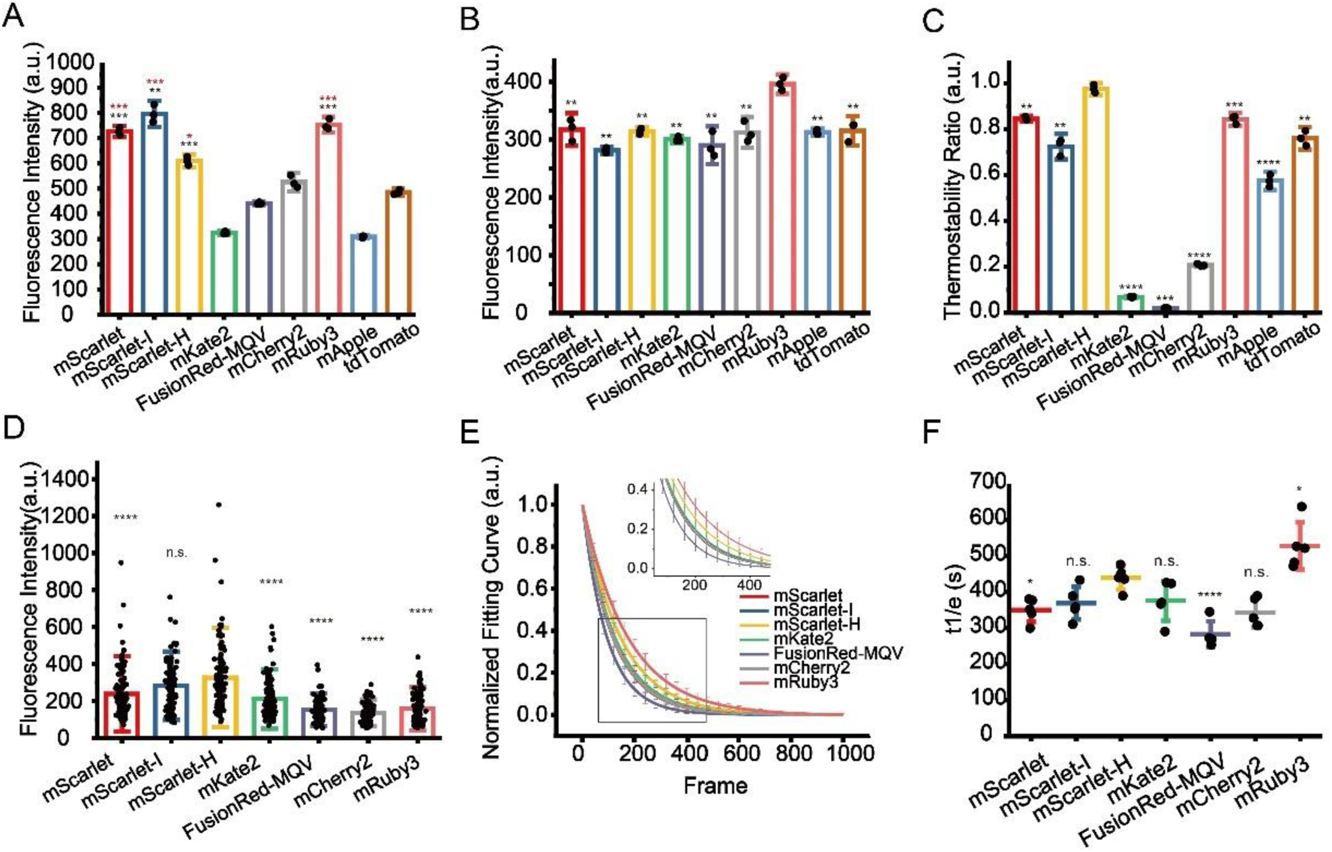
Identification of optimal RFPs for CLEM imaging. (**A**) Residual fluorescence intensity of RFPs after 1% OsO4 post-fixation. Red stars represent comparisons to mCherry2, black stars represent comparisons to mKate2. (**B**) Residual fluorescence intensity of RFPs after 1% OsO4 post-fixation followed by absolute ethanol dehydration. (**C**) Thermostability of RFPs at 60 °C. (**D**) Fluorescence intensity of RFPs on Epon-embedded sections (100 nm). (**E**) Normalized photobleaching curves of RFPs on Epon-embedded sections (100 nm). Enlarged view of the boxed area is shown on top. (**F**) Statistics of the photobleaching time when the fluorescence intensity of RFPs reduced to 1/e of their initials. Bars represent mean ± SD. P values were determined with two-tailed t-tests in (**A-C**) (n = 3) and (**F**) (n = 5), Mann-Whitney U test was performed in (**D**) (n = 106). n.s. indicates p > 0.05, * indicates p < 0.05, ** indicates p < 0.01, *** indicates p < 0.001, **** indicates p < 0.0001. Data are summarized in Table S5 & 9.

Next, we performed an *in vitro* thermostability test at 60 °C to mimic the effect of high-temperature on fluorescence preservation during Epon resin polymerization. Among nine selected RFPs, mScarlet-H showed the best thermostability that is significantly higher compared to the others (Figure 2C). After complete EM sample preparation, mScarlet-H showed the highest residue fluorescence, followed by mScarlet-I, which are both significantly higher than the previously reported mKate2 and mCherry2, while the residue fluorescence of mRuby3 was surprisingly low (Figure 2D). Statistic results showed that mScarlet-H has the highest SBR among tested RFPs (Figure S7). We speculated that mRuby3 might be super sensitive to the Epon embedding procedure. In addition, even though mApple and tdTomato show comparable fluorescence residue to mKate2 and mCherry2 after OsO_4_ fixation and dehydration treatment, and even better thermostability than mCherry2 and mKate2, they totally lose their fluorescence after complete EM sample preparation (data not shown).

Additionally, we also measured the photostability of these RFPs on standard EM sample sections under the illumination condition of CLEM imaging. mScarlet-H had higher photostability than other RFPs except mRuby3 (Figure 2E, F).

Taken together, we recommend mScarlet-H as the best red probe for standard CLEM as it has the highest in-resin fluorescence and a preferable photostability among RFPs after complete TEM sample preparation.

### mScarlet-H is a generalizable red probe for post-Epon-embedding CLEM

To exemplify the utility of mScarlet-H in Epon-embedded same-section CLEM, we constructed mScarlet-H fusions with the mitochondria targeting peptide, the endoplasmic reticulum membrane protein Sec61β, and histone H2B, individually. Cell sections transiently expressing mScarlet-H fusions were prepared according to the standard TEM sample preparation procedure. The fluorescence signal of mScarlet-H from thin sections (100 nm) was consecutively recorded for multiple frames. The first 100 frames were averaged to smooth the background noise and increase the signal-to-noise ratio. Mitochondria, endoplasmic reticulum, and Histone H2B were correctly targeted and clearly identified under fluorescence microscopy (FM) (Figure 3A, D, G). Next, to obtain the corresponding ultrastructure of these cell components, the same sections were subsequently imaged using TEM (Figure 3B, E, H). All three structures were successfully aligned with high accuracy (Figure 3C, F, I). These results demonstrate that mScarlet-H is a high-performance red probe that can be generalized for same-section CLEM imaging of various cellular targets.

**Figure 3.**
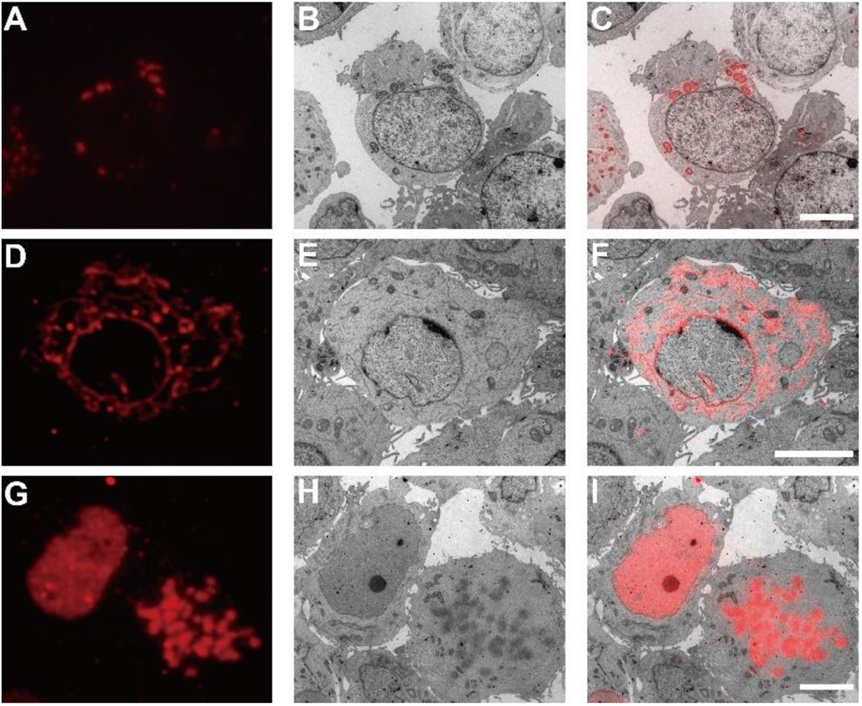
Post-Epon-embedding CLEM imaging by mScarlet-H. FM, EM, and CLEM images of the mitochondrial matrix (**A-C**), the endoplasmic reticulum sec61β protein (**D-F**), and the nucleosome H2B protein (**G-I**) labeled by mScarlet-H (100 nm slices). Scale bars, 5 µm.

### Dual-color post-Epon-embedding same-section CLEM using mEosEM-E and mScarlet-H

mScarlet-H and mEosEM-E were then combined to perform dual-color same-section CLEM imaging. The nuclear envelope (laminA) and mitochondria matrix in HEK 293T cells were labeled by mScarlet-H and mEosEM-E, respectively. After standard EM sample preparation, dual-color wide-filed FM images of cell sections (100 nm) were sequentially acquired with a 561-nm laser for 100 frames to image mScarlet-H, followed by 20 frames with a 488-nm laser, and 20 frames with 488-plus 405-nm lasers to image mEosEM-E. For the red channel, images were averaged to obtain the final image; while for the green channel, sLM was applied as mentioned above. As shown in Figure 4A, the mScarlet-H-labeled laminA in the red channel exhibited a good nuclear envelope structure with a high SBR. However, in the green channel, regardless of whether mEosEM-E was in the on- (Figure 4B) or off-state (Figure 4C), the SBR of the fluorescence signal was very low. Furthermore, in addition to mEosEM-E-labeled mitochondrial structure, nuclear envelope structure due to mScarlet-H crosstalk could also be observed as indicated by the merged image of mEosEM-E (on-state) and mScarlet-H (Figure 4D, box). Notably, using the photoswitching property of mEosEM-E and the sCLEM method, the structure of mitochondria could be obtained with a very high SBR (Figure 4E). Moreover, the mScarlet-H crosstalk signal was effectively eliminated (Figure 4F) by the sCLEM imaging scheme, as the green signal of mScarlet-H does not possess the photoswitching property. A further investigation showed that the crosstalk phenomenon is universal for all the tested RFPs that survived standard EM sample preparation (Figure S8). A very likely reason is that under high-energy illumination, severe chromophore photoreduction happens that leads to obvious red-to-green photoconversion of the RPFs, a mechanism that has been reported before [24–26].

**Figure 4.**
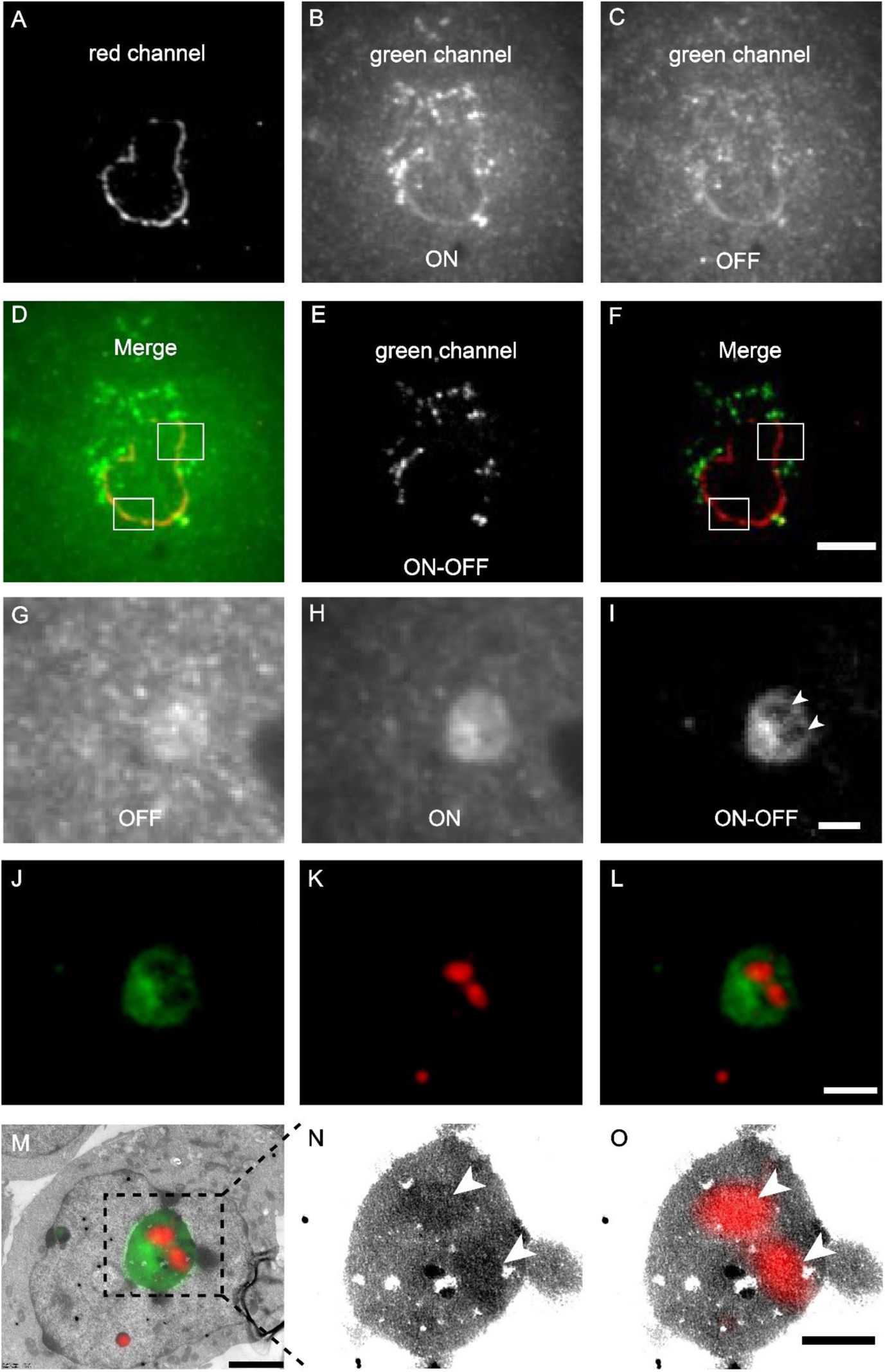
Dual-color post-Epon-embedding CLEM using mEosEM-E and mScarlet-H. (**A-F**) Dual-color imaging of mEosEM-E-labeled mitochondria and mScarlet-H-labeled LaminA protein in HEK 293T cell sections (100 nm). Representative red channel (**A**), green channel ON (**B**), and OFF (**C**) FM images. (**D**) Merged image of (**A**) and (**B**). White boxes indicate areas that showed crosstalk signals from mScarlet-H. (**E**) Green channel ON ─ OFF image. (**F**) Merged image of (**A**) and (**E**). White boxes indicate the same areas in (**D**). Scale bar, 5 µm. (**G-O**) Dual-color CLEM imaging of nucleolar proteins. OFF (**G**), ON (**H**), and ON ─ OFF (**I**) images of mEosEM-E labeled B23 in HEK 293T cells sections (100 nm). White arrowheads indicate areas that were not visible in (**G**) and (**H**). Scale bar, 2 µm. Green channel FM (**J**), Red channel FM (**K**), merged channel FM (**L**), and CLEM (**M**) images of HEK 293T cell sections expressing mEosEM-E labeled B23 and mScarlet-H labeled Nopp140. Gamma value: 1.6 for both channels. Scale bars, 2 µm. (**N-O**) Enlarged EM (**N**) and CLEM (**O**) images of boxed area in (**M**), white arrowheads indicate the localization of Nopp140 that showed higher EM contrast compared to B23. Gamma value: 1.6 for red channel. Scale bar, 1 µm.

Next, we applied dual-color CLEM to investigate how nucleolar proteins are organized in the nucleolus. The nucleolus of mammalian cells has three morphologically distinct components: the fibrillar center (FC), the dense fibrillar component (DFC), and the granular component (GC), which perform different functions in ribosome biogenesis [27]. We labeled the GC marker B23 with mEosEM-E and labeled Nopp140, whose localization has been controversially reported to be at the FC or DFC [28, 29], with mScarlet-H. The results showed that in the green channel, typical GC structures were clearly visible in FM images after sLM, again exemplifying the utility of the subtraction strategy (Figure 4G-I). Merging of green and red channels revealed that the red signal was encircled by the green signal (Figure 4J-L).

Further correlative FM and EM images (Figure 4M-O) showed that the fluorescence signal of Nopp140 was closely matched to the solid high electron-dense regions (Figure 4N, arrowheads) in the EM image, indicating a DFC localization, which is consistent with the fact that it’s a functional different area other than GC (Figure 4M, O). Additionally, we also double-labeled Nop52 and Nopp140 by mEosEM-E and mScarlet-H, respectively. Unexpectedly, while Nop52 was often used as a marker of GC [30], Nopp140 was not located in the vacant holes devoid of the Nop52 signals, but partially overlapped with Nop52 (Figure S9). These results indicated that Nop52 localized both at the GC and the DFC.

Taken together, these results strongly demonstrate that mEosEM-E and mScarlet-H are optimal probes for high-quality dual-color post-Epon-embedding CLEM imaging, and sCLEM with mEoEM-E offers a great advantage in reducing the green background of the Epon resin as well as in eliminating the signal crosstalk of RFPs.

## Discussion

In CLEM imaging, there is always a trade-off between the performance of the FM and the EM. Thus, it is challenging to obtain good FM and EM images of the same ultrathin section at the same time. Developing CLEM probes with resistance to the treatment of EM sample preparation is an efficient way to solve this problem, thus, making CLEM with both high-quality FM and EM possible.

Post-Epon-embedding CLEM was enabled by several FPs reported very recently, however, when these probes were used in practical CLEM imaging, the SBR of FM is very low. The problem is more prominent for imaging low-abundance proteins. In the current study, we utilized the optical switching properties of RSFPs and proposed an sCLEM method, by which the FP signal can be easily distinguished from that of the unmodulated background. sCLEM works more efficiently than the OLID technique in our case, which may be due to the phenomenon that the background fluorescence also showed minor on and off responses to 405- and 488-nm excitation. Compared to the single-frame subtraction (Max-Min), subtraction between the multi-frame averages of the on and off states (Sum_AVG_-Sum_AVG_) has an additional benefit of largely attenuated noise. Notably, the sCLEM method hits two birds with one stone. In addition to effectively removing the background signal of the Epon resin and improving the SBR of the FM image, it also removes the signal crosstalk of the RFP in the green channel during dual-color CLEM. To further improve the SBR of FM image, we used mEosEM as the template and developed a variant mEosEM-E with enhanced on-state brightness and on/off contrast that is beneficial for fluorescent background elimination and weak signal extraction. We also identified mScarlet-H as the best RFP for post-Epon-embedding CLEM. This was validated by actual imaging applications, where we proved that mEosEM-E/mScarlet-H is an excellent FP pair for same-section dual-color CLEM of post-Epon-embedded samples. Compared with mEos4b, mEosEM retained higher fluorescence intensity after osmic acid treatment [16]. Considering that the fluorescence intensity retained in GMA-embedded samples is mainly dependent on the osmic acid-resistant properties of the FP, we suspected that mEosEM-E/mScarlet-H would also be excellent choices for the hydrophilic GMA resin.

Besides the development of an advanced green CLEM probe, we also performed a meticulous study that thoroughly dissected the influence of key chemical treatments during TEM sample preparation on the fluorescence preservation of RPFs. Interestingly, we found that the responses of RFPs to these treatments vary greatly. Some RFPs (mRuby3, tdTomato) showed good O_S_O_4_ resistance, however, their superiorities diminished after complete EM sample preparation, suggesting that they are more sensitive to the subsequent sample treatments. Therefore, we reasoned that the O_S_O_4_ resistance assay is not a suitable gold standard for CLEM probe screening, even though OsO_4_ fixation seems to be the uppermost factor for fluorescence reduction. Development of an optimal probe requires investigation of the probe’s final performance after complete sample preparation, even though this will significantly increase the workload of probe development, and add challenges for high-throughput screening. Furthermore, this study also provided useful information that may assist the rational design of new types of probes. For example, mRuby3 exhibited superior robustness towards dehydration, which may provide clues for developing FPs that can work in a hydrophobic environment.

Before this study, nucleolar proteins have never been inspected by the same section CLEM. Our dual-color CLEM results revealed that although both have been considered as a GC marker, Nop52 and B23 have different sub-nucleolar localizations. The partial co-localization of Nop52 with Nopp140 suggests that B23 would be a better marker to represent GC. Moreover, we confirmed a DFC localization of Nopp140 in our experimental conditions.

Further directions for the development of post-Epon-embedding CLEM probes would be: 1) the establishment of a high-throughput automatic on-section screening system; 2) a red RSFP CLEM probe for background-reduced and/or SR dual-color CLEM, for which mScarlet-H may be a good starting template; 3) both green and red CLEM probes with even higher on-section brightness, SBR, and photostability.

## Materials and methods

### Development of mEosEM-E

Saturation mutagenesis of mEosEM at His63 was performed in the pEGFP-N1-mito-mEosEM plasmid with Q5 high fidelity polymerase (New England Biolabs). The amplified fragments containing homologous arms and mutation sites were transformed into the Top10 competent cells (Tsingke) and sequenced (Tsingke). HEK 293T cells were transfected with sequencing validated mutants and were prepared following the standard EM sample preparation procedure. Cell slices with a thickness of 100 nm were imaged under a homemade wide field fluorescence microscope (Olympus IX71 body and Olympus PLAN APO 100×, 1.49 NA oil objective) to characterize the photoswitching properties of different mutants. mEosEM-E was identified for its highest photoswitching contrast ratio.

### Plasmid construction

For prokaryotic expression plasmids pRsetA-RFPs (mScarlet, mScarlet-I, mScarlet-H, mKate2, FusionRed-MQV, mCherry2, mRuby3, mApple, tdTomato), RFPs fragments were PCR amplified and digested with BamHI and EcoRI restriction enzymes. Then fragments were ligated into the pRsetA vector digested with the same enzymes. For eukaryotic expression plasmids, pmEosEM-N1-mito was digested with AgeI and NotI restriction enzymes to replace mEosEM with mEosEM-E or RFPs fragments with the same restriction enzyme sticky ends. pEGFP-C1-Sec61, pmEos3.2-N1-H2B, and pmEosEM-C1-LaminA plasmids were digested with AgeI/BglII, XhoI/NotI, and NheI/BglII restriction enzymes respectively to replace EGFP, mEos3.2 and mEosEM with Scarlett-H fragment. For the construction of pmEosEM-C1-B23, pmEosEM-C1-Nop52, and pmScarlet-H-C1-Nopp140 plasmids, the full-length cDNA of B23, Nop52 and Nopp140 were PCR amplified from the HEK 293T cDNA library, digested with EcoRI/SalI, HindIII/SalI, and BglII/SalI restriction enzymes and inserted into pEGFP-C1-mEosEM-LaminA and pEGFP-C1-mScarlet-H-LaminA plasmids to replace LaminA. Q5 polymerase and T4 ligase were purchased from New England Biolabs. All restriction enzymes were purchased from Thermo Fisher Scientific.

### Cell culture and transfection

HEK 293T and U-2 OS cells were cultured in Dulbecco’s Modified Eagle Medium (Gibco) and McCoy’s 5A Modified Medium (Gibco), respectively, supplemented with 10% FBS (Gibco) and 1% penicillin-streptomycin (TransGene Biotech). Cells were maintained at 37°C in an incubator supplied with 5% CO2 (vol/vol). Transfections were performed with purified plasmids using lipofectamine 2000 (Invitrogen) (with a ratio of 1 µg DNA: 3 µl lipofectamine) in 6 cm pertri dishes or 12-well cell culture plates following the manufacturer’s instructions.

### EM sample preparation

Successfully transfected cells were trypsinized and then harvested by centrifugation. Cells were fixed with 2% paraformaldehyde (Electron Microscopy Sciences) and 2.5% glutaraldehyde (Electron Microscopy Sciences) in 100 mM PBS at 4°C overnight. Pre-fixed cells were washed 2 times with PBS buffer and 2 times with Milli-Q water on ice. Then post-fixation with 1% OsO_4_ was performed on ice for 1 h. After 4 washes with Milli-Q water, cells were gradually dehydrated with a series of concentration gradients of ethanol (30% 50% 70% 90% 100%) for 6 min each and followed by dehydration with 100% acetone for 6 min. Cells were then infiltrated step by step in mixtures of Epon (50% Epon812, 30.5% NMA, 18% DDSA, 1.5% DMP-30) and acetone with gradient concentrations (50% Epon for 2 h, 75% Epon for 3 h, 100% Epon for 12 h, 100% Epon for 12 h). Finally, cells were embedded in 100% Epon at 60°C for 16 h and then sectioned into 100-nm slices for subsequent experiments.

### Photoswitching property analysis

HEK 293T cells expressing mitochondria targeted mEosEM and mutants were used for photoswitching property analysis. For the analysis before EM sample preparation, cells were excited by continuous 488-nm laser (0.41 kW/cm_2_), while every 10 s, a 405-nm laser (0.21 kW/cm_2_) pulse of 0.1 s were added to turn on the FPs. Six cycles were recorded for all FPs. For analysis after EM sample preparation, 100-nm cell sections were imaged under a continuous 488-nm laser for 50 frames, after which the 405-nm laser was added for 1 s to record the fluorescence signal of the FPs at the on-state. The contrast ratio was calculated as follow:

Contrast ratio = (Max-Mean)/Mean

Max represents the maximum value of the signal in all acquired images, Mean represents the averaged signal value of the first 20 frames acquired before the application of 405-nm laser.

### High-content analysis of pre-fixation, post-fixation, and dehydration resistance of RFPs

U-2 OS cells expressing pRFPs-mito were seeded into 96-well optical polymer base microplates and fixed with 2% paraformaldehyde (Electron Microscopy Sciences) and 2.5% glutaraldehyde (Electron Microscopy Sciences) in 100 mM PBS at 37°C for 15min. The pre-fixed cells were washed 4 times with PBS and then post fixed with 1% OsO_4_. After 10 min incubation at 4°C, OsO_4_ was removed and cells were washed 5 times with PBS and stored in PBS. For dehydration, PBS buffer was replaced with absolute ethanol for 20 min without washing. Fluorescence images were acquired by the Opera Phenix_TM_ High Content Screening System (PerkinElmer) using a 20×, 0.4 NA water objective with an excitation laser of 568 nm. Data quantification and analysis were performed using Harmony 4.9 software (PerkinElmer).

### Protein purification and thermostability measurement

BL21(DE3) competent cells (Tsingke) were transformed with prokaryotic expression plasmids pRsetA-RFPs and single clones were cultured in liquid LB medium to the logarithmic growth phase. Then 0.8 mM IPTG (Isopropyl β-d-1-thiogalactopyranoside) was added to induce the expression of RFPs. After induction at 16 °C for 24 h, cells were harvested by centrifugation. The pelleted bacteria were resuspended in binding buffer including 10 mM imidazole and lysed by ultrasonication. Protein was purified through affinity chromatography (Ni-NTA His-Bind resin, Qiagen), followed by gel filtration chromatography (Superdex 200 Increase 10/300 GL, GE Healthcare). Purified protein was diluted in PBS buffer (pH = 7.2) and the fluorescence intensity was recorded at 60 °C in the Rotor-Gene 6600 real-time PCR cycler (Qiagen) for 18 h. The thermostability was defined as the ratio between the initial and the final fluorescence intensities.

### Photostability measurement

HEK 293T cell samples expressing mitochondria targeted RFPs were prepared as described above for EM and sectioned into 100 nm slices. Sample slices were illuminated with a 561-nm laser and the fluorescence signal was acquired by time-lapse imaging. Photostability was defined as the time when fluorescence intensity reached 1/e of its initial.

### CLEM imaging

Cleaning and coating of the coverslips with pioloform (Ted Pella) were processed as previously reported [31]. Sectioned slices were placed on well prepared coverslips and submerged with mowiol buffer to recover the fluorescence. After a 30 min incubation, slices were imaged using a widefield fluorescence microscope (Olympus IX71) equipped with a 100×, 1.49 NA oil objective (Olympus PLAN APO). Images were acquired using an electron-multiplying charge-coupled device (EMCCD) camera (Andor iXon DV-897 BV). For the red channel, 100 frames were acquired for averaging during excitation by a 561-nm laser. For the green channel, 20 frames were first acquired using 488-nm laser excitation alone, then another 20 frames were acquired while a 405-nm laser pulse of 1 s was added to switch on the mEosEM-E molecules. For both channels, the exposure time was 50 ms. After fluorescence signal recording, DIC images of 100×, 16×, and 10× magnifications were sequentially collected to assist the target retrieving during subsequent EM imaging. After LM imaging, a rectangle on the pioloform film with a slice on it was scored by a knife. 12% hydrofluoric acid was dropped on the periphery of the rectangle to detach the pioloform film from the glass coverslip. When the coverslip was submerged under water, the detached pioloform film would float on the surface and was captured by an uncoated slot grid. Next, the section slice was stained with 2% UA and 1% Sato’s triple lead. Finally, the sample was imaged under TECNAI SPIRIT TEM (FEI). Gold nanoparticles were used as the fiducial marker. FM and EM images were correlated by eC-CLEM following the previous protocol [16].

### Registration of green and red channel FM images

TetraSpeck™ Microspheres (Thermofisher Scientific) were diluted in PBS buffer and spotted on a coverslip (Fisher Scientific). Dual-color fluorescence signals were recorded simultaneously under 488- and 561-nm lasers. The registration was performed with the Fiji plugin “Multi Registration” in ImageJ.

## Acknowledgments

We are grateful to Ya Wang from Research Platform for Protein Sciences at Institute of Biophysics for her support of Opera Phenix_TM_ High Content Screening system data analysis. We thank Center for Biological Imaging at Institute of Biophysics for EM imaging support.

## Author contributions

P.X., M.Z., D.P., and N.L designed the experiments. D.P., N.L., and W.H. performed the experiments. P.X., M.Z., D.P., and N.L. analyzed the data. P.X., M.Z., and T.X. supervised the project and wrote the manuscript. K.D provided guidance on the project.

## Supplementary Materials

|  |  |
| --- | --- |
| **Figure S1** | Autofluorescence of Epon resin slice |
| **Figure S2** | Photoswitching properties of mEosEM and mutants before EM sample preparation |
| **Figure S3** | Scheme of sLM imaging of EM sample |
| **Figure S4** | SBR comparison of mEosEM and mEosEM-E in OFF- and ON-images |
| **Figure S5** | Pre-fixation influence on RFPs |
| **Figure S6** | Fluorescence intensity of RFPs in pre-fixed cells followed by dehydration |
| **Figure S7** | SBR comparison of RFPs after EM sample preparation |
| **Figure S8** | Green fluorescence emission properties of RFPs |
| **Figure S9** | Dual-color post-Epon-embedding CLEM imaging of Nopp52 and Nopp140 |
| **Table S1** | Statistics for contrast ratio comparison between mEosEM and mEosEM-E |
| **Table S2** | Statistics for SBR of mEosEM in OFF, ON, and ON — OFF images |
| **Table S3** | Statistics for SBR of mEosEM-E in OFF, ON, and ON — OFF images |
| **Table S4** | Statistics for SBR comparison between mEosEM and mEosEM-E in ON — OFF images |
| **Table S5** | Statistics for fluorescence intensity comparison of RFPs after 1% OsO<sub>4</sub> post-fixation |
| **Table S6** | Statistics for fluorescence intensity comparison of RFPs after 1% OsO<sub>4</sub> post-fixation and dehydration treatment |
| **Table S7** | Statistics for thermostability comparison of RFPs |
| **Table S8** | Statistics for fluorescence intensity comparison of RFPs after EM sample preparation |
| **Table S9** | Statistics for photostability comparison of RFPs |
| **Table S10** | Statistics for SBR comparison between mEosEM and mEosEM-E in OFF images |
| **Table S11** | Statistics for SBR comparison between mEosEM and mEosEM-E in ON images |
| **Table S12** | Statistics for fluorescence intensity comparison of RFPs in live cells |
| **Table S13** | Statistics for fluorescence intensity comparison of RFPs after pre-fixation |
| **Table S14** | Statistics for fluorescence intensity comparison of RFPs after dehydration treatment |
| **Table S15** | Statistics for SBR comparison of RFPs after EM sample preparation |

**Figure S1.**
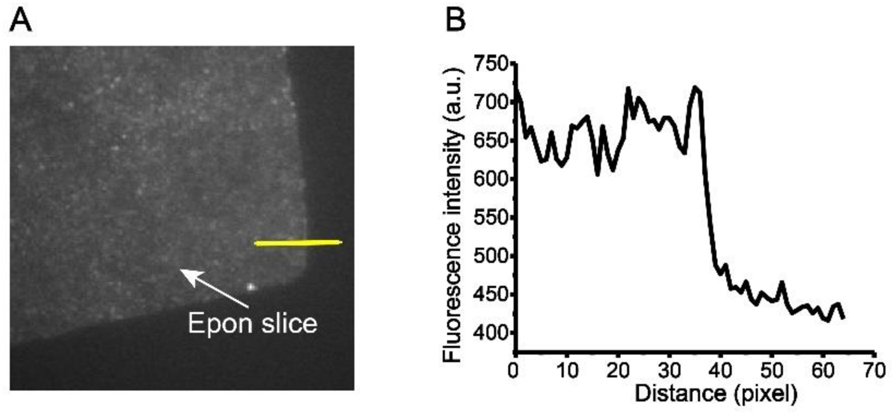
Autofluorescence of Epon resin slice. (**A**) A corner of Epon slice was imaged under 488-nm laser (0.41 kW/cm_2_). (**B**) Fluorescence intensity profile of the yellow line in (**A**). Fluorescence intensity was plotted against distance.

**Figure S2.**
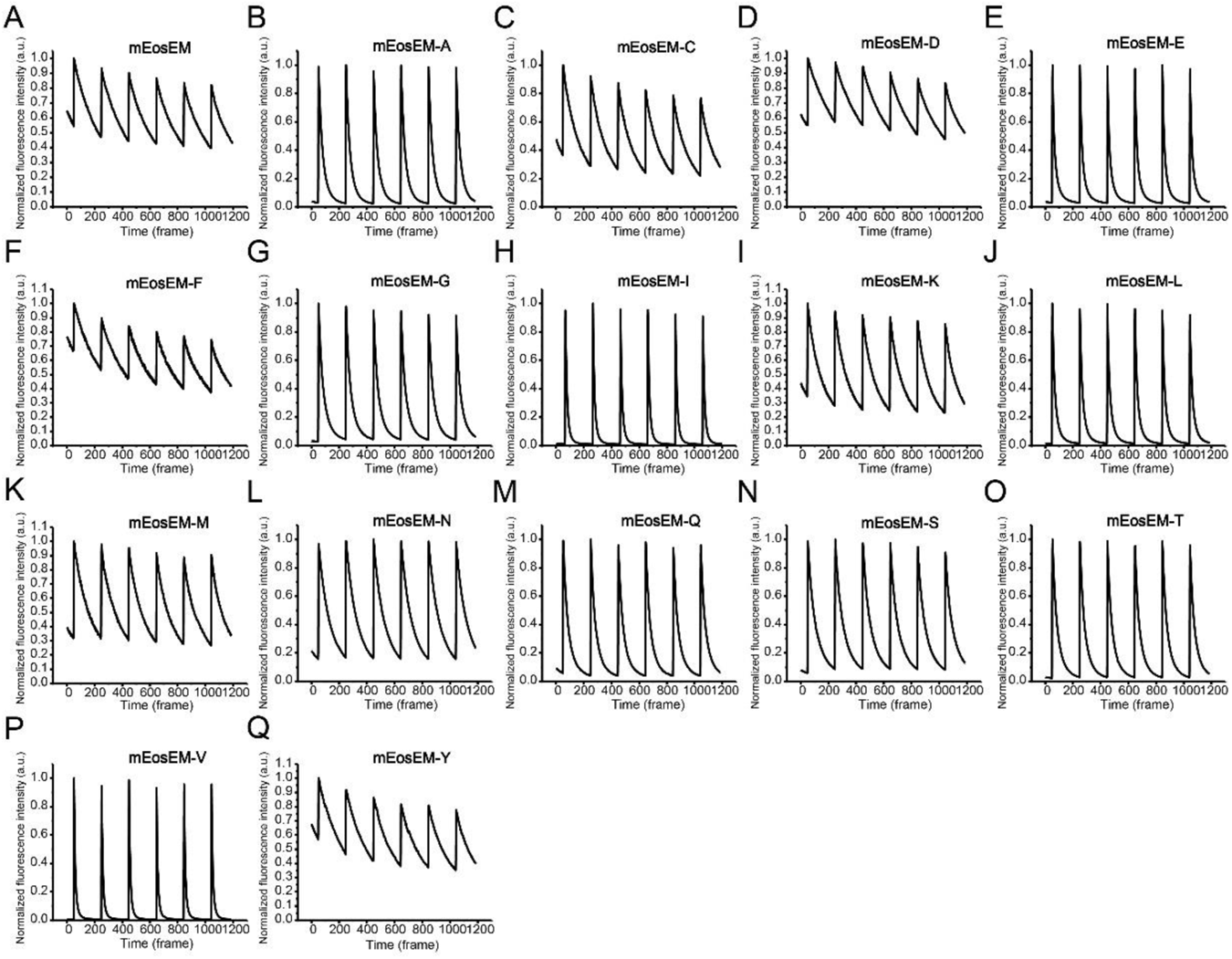
Photoswitching properties of mEosEM and mutants before EM sample preparation. (**A-Q**) Normalized fluorescence intensity of mEosEM and mutants was plotted against time. Cells expressing different fluorescent proteins were continuously illuminated with a 488-nm laser (8 W/cm_2_), while every 10 s, a 405-nm laser pulse (0.1 s, 9 W/cm_2_) were applied to switch on the FPs. Exposure time, 50 ms.

**Figure S3.**
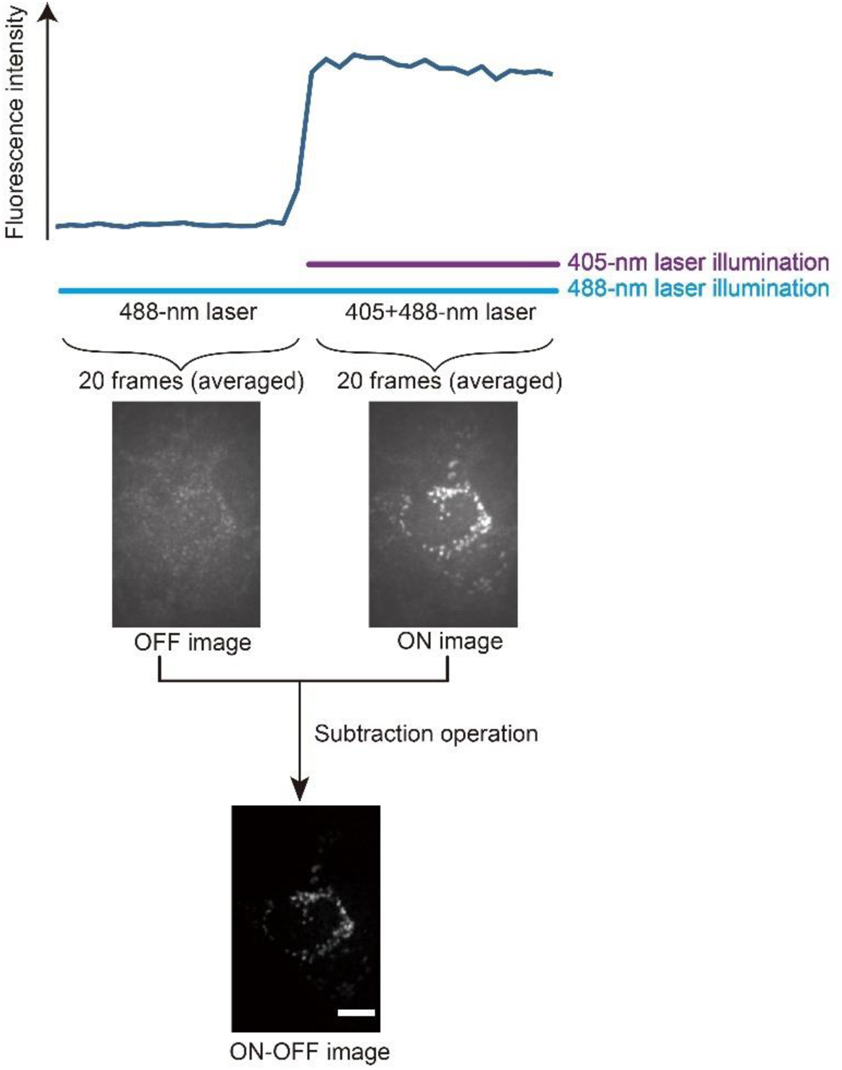
Scheme of sLM imaging of EM sample. 100-nm cell sections were imaged under a continuous 488-nm laser for 50 frames, after which the 405-nm laser was added for 20 frames to record the fluorescence signal of the FPs at the on-state. The 20 frames immediately before 405-nm illumination were averaged to produce OFF images, and the subsequent 20 frames with 405-nm illumination were averaged to produce ON images. The sLM images (ON ─ OFF images) were generated by pixel-by-pixel subtraction using adjacent ON and OFF images. Scale bar, 5µm.

**Figure S4.**
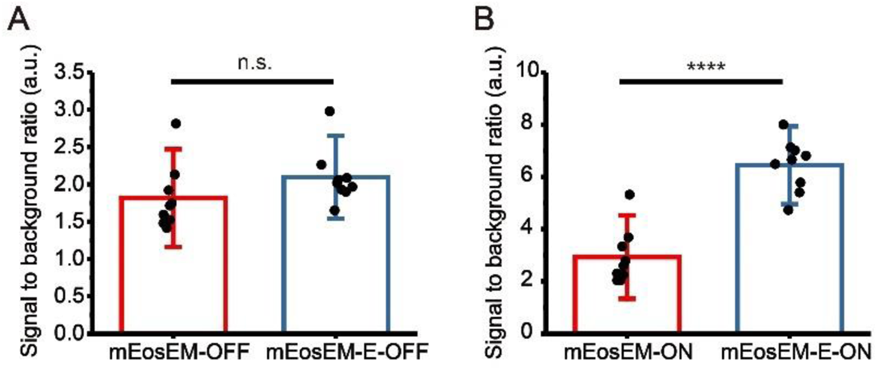
SBR comparison of mEosEM and mEosEM-E in OFF- and ON-images. (**A**) Statistics of SBR in OFF-images between mEosEM and mEosEM-E. (**B**) Statistics of SBR in ON-images between mEosEM and mEosEM-E. Bars represent mean ± SD. P-value were determined with two-tailed t-test in (**A-B**) (n = 9). n.s. indicates p>0.05, **** indicates p<0.0001. Data are summarized in Table S10 & 11.

**Figure S5.**
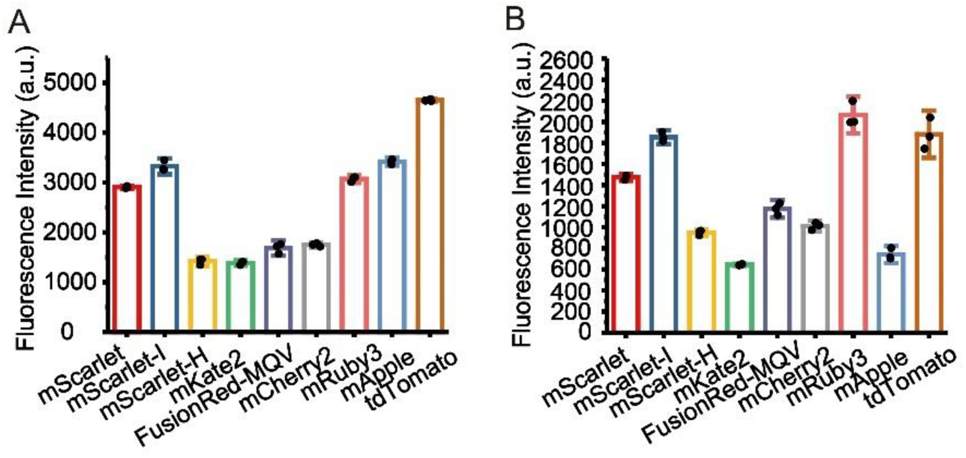
Pre-fixation influence on RFPs. (**A**) Fluorescence intensity of RFPs in live cells. Cells expressing RFPs were seeded in 96-well plates and imaged. (**B**) Fluorescence intensity of RFPs in pre-fixed cells. Cells expressing RFPs were seeded in 96-well plates and treated with fixation buffer (2.5% Glutaraldehyde and 2% Paraformaldehyde in PBS) for 15 min then washed 3 times with PBS buffer. Data were recorded using a high-content screening system. Excitation wavelength, 561 nm. Exposure time, 40 ms. Bars represent mean ± SD (n = 3). Data are summarized in Table S12 & 13.

**Figure S6.**
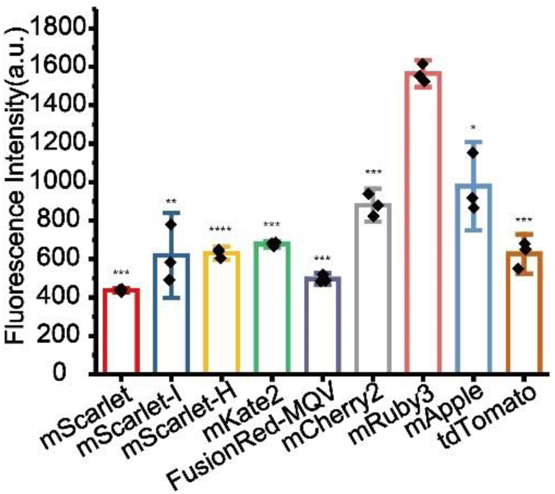
Fluorescence intensity of RFPs in pre-fixed cells followed by dehydration. Cells expressing RFPs were seeded in 96-well plates and fixed with fixation buffer (2.5% Glutaraldehyde and 2% Paraformaldehyde in PBS buffer) for 15 min, and then treated with absolute ethanol for 20 min. Data were recorded using a high-content screening system. Excitation wavelength, 561 nm. Exposure time, 40 ms. Bars represent mean ± SD. P-value were determined with two-tailed t-tests (n = 3). * indicates p<0.05. ** indicates p<0.01. *** indicates p<0.001. **** indicates p<0.0001. Data are summarized in Table S14.

**Figure S7.**
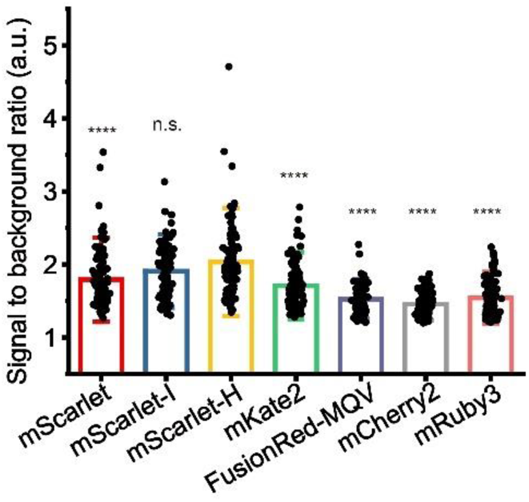
SBR comparison of RFPs after EM sample preparation. HEK 293T cells expressing RFPs were prepared under standard EM sample preparation and sectioned into 100 nm slices. Mann-Whitney U test was performed (n = 106). n.s. indicates p > 0.05, **** indicates p < 0.0001. Data are summarized in Table S15.

**Figure S8.**
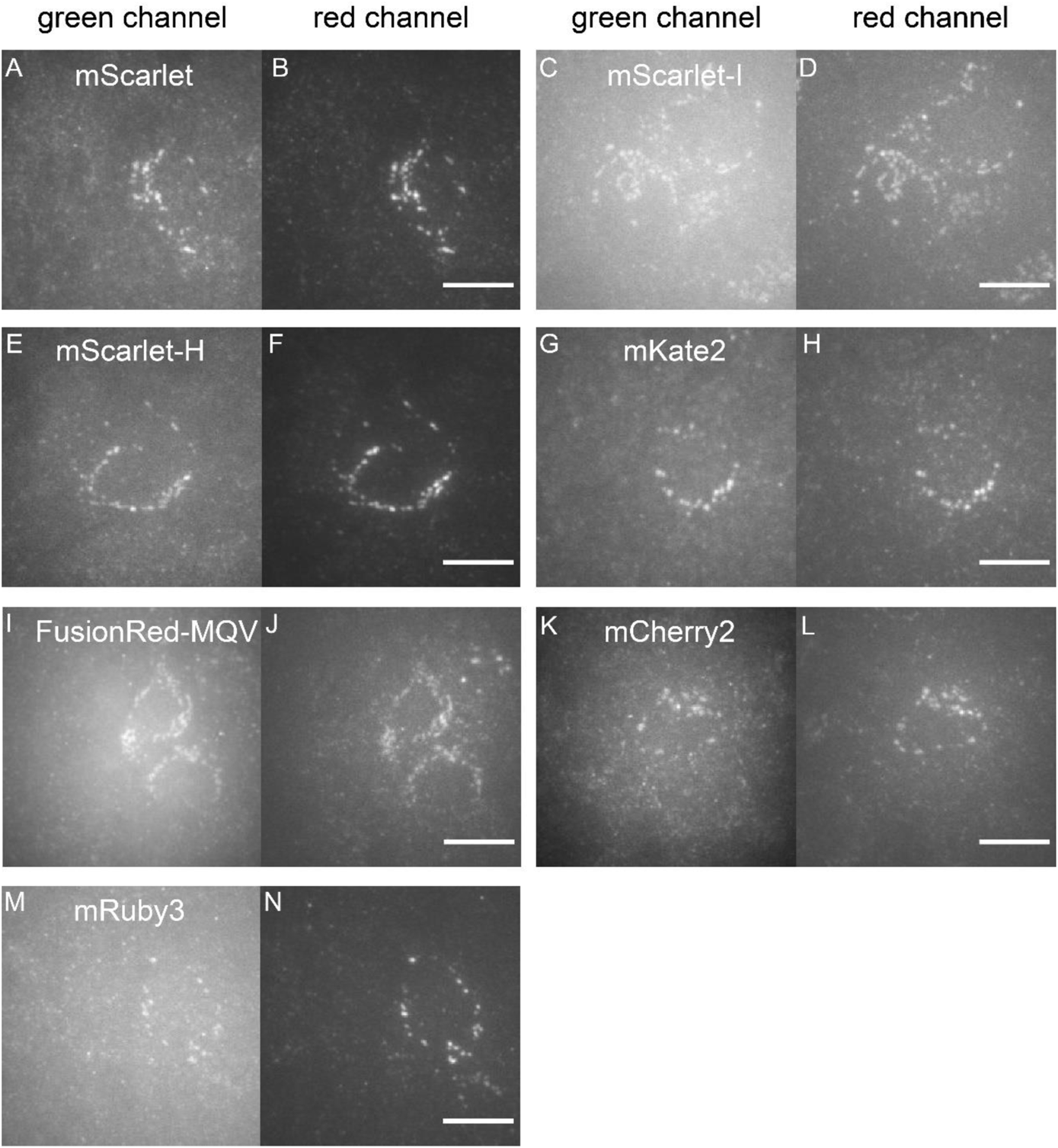
Green fluorescence emission properties of RFPs. HEK 293T cells expressing RFPs were prepared under standard EM sample preparation and sectioned into 100 nm slices. Sample slices were imaged by wide field fluorescence microscopy with sequential illumination of 561- and 488-nm laser. (**A**), (**C**), (**E**), (**G**), (**I**), (**K**) and (**M**) Slices were illuminated under 488-nm laser (0.92 kW/cm_2_) and signals were recorded from the green channel. (**B**), (**D**), (**F**), (**H**), (**J**), (**L**) and (**N**) Slices were illuminated under 561-nm laser (0.57 kW/cm_2_) and signals were recorded from the red channel. Scale bars, 10 μm.

**Figure S9.**
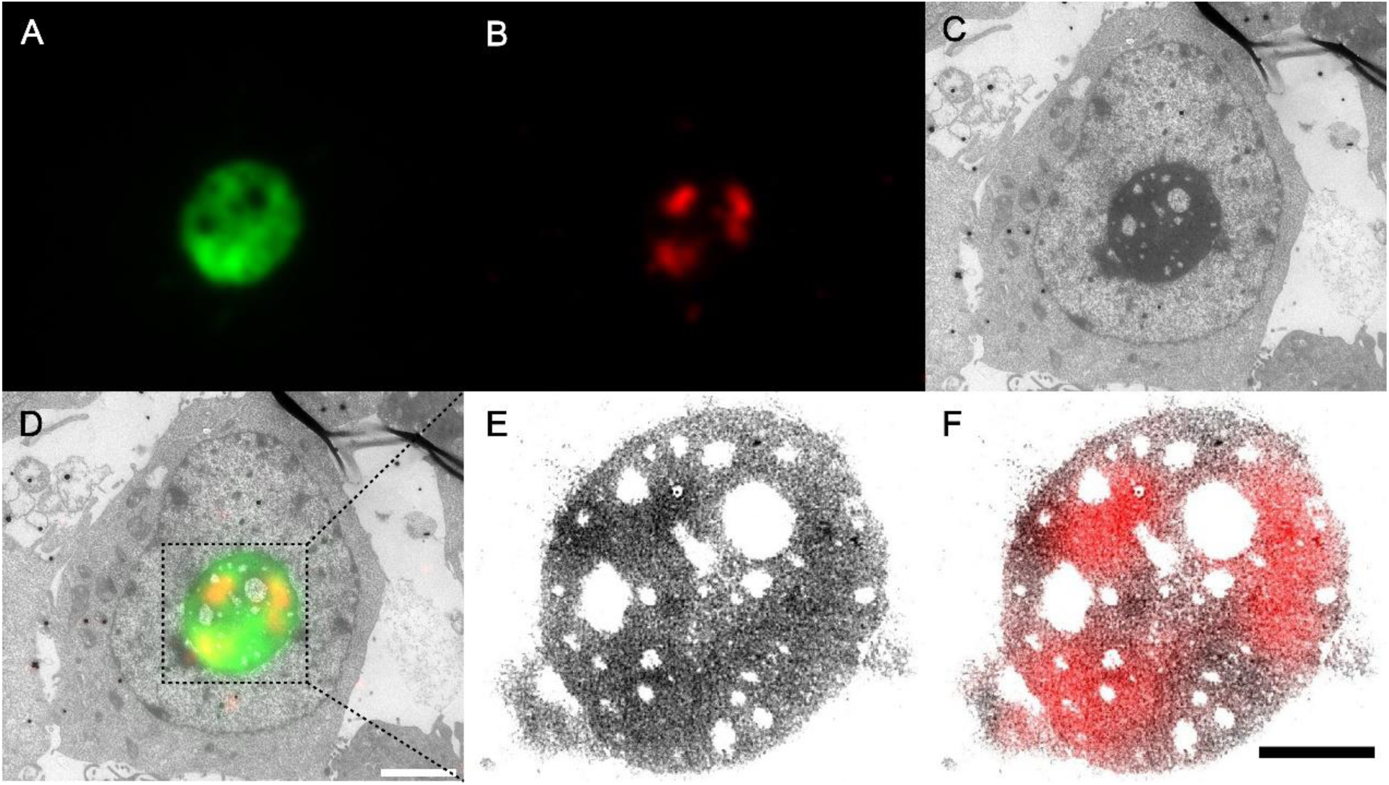
Dual-color post-Epon-embedding CLEM imaging of Nopp52 and Nopp140. (**A-D**) Dual-color CLEM imaging of mEosEM-E labeled Nopp52 and mScarlet-H labeled Nopp140 protein in HEK 293T cell sections (100 nm). Green channel FM (**A**), red channel FM (**B**), EM (**C**), and CLEM (**D**) images. Scale bar, 2 µm. (**E-F**) Enlarged EM (**E**) and red channel CLEM (**F**) images of boxed area in (D). Gamma value: 1.6 for both channels. Scale bar, 1 µm.

**Table S1.**
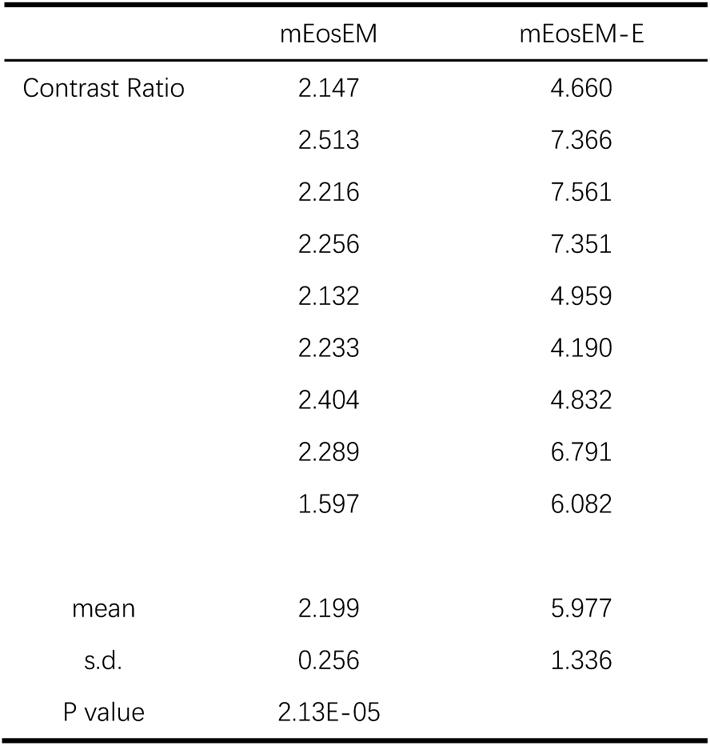
Statistics for contrast ratio comparison between mEosEM and mEosEM-E. Two-tailed t-test was performed between mEosEM and mEosEM-E, n = 9.

**Table S2.**
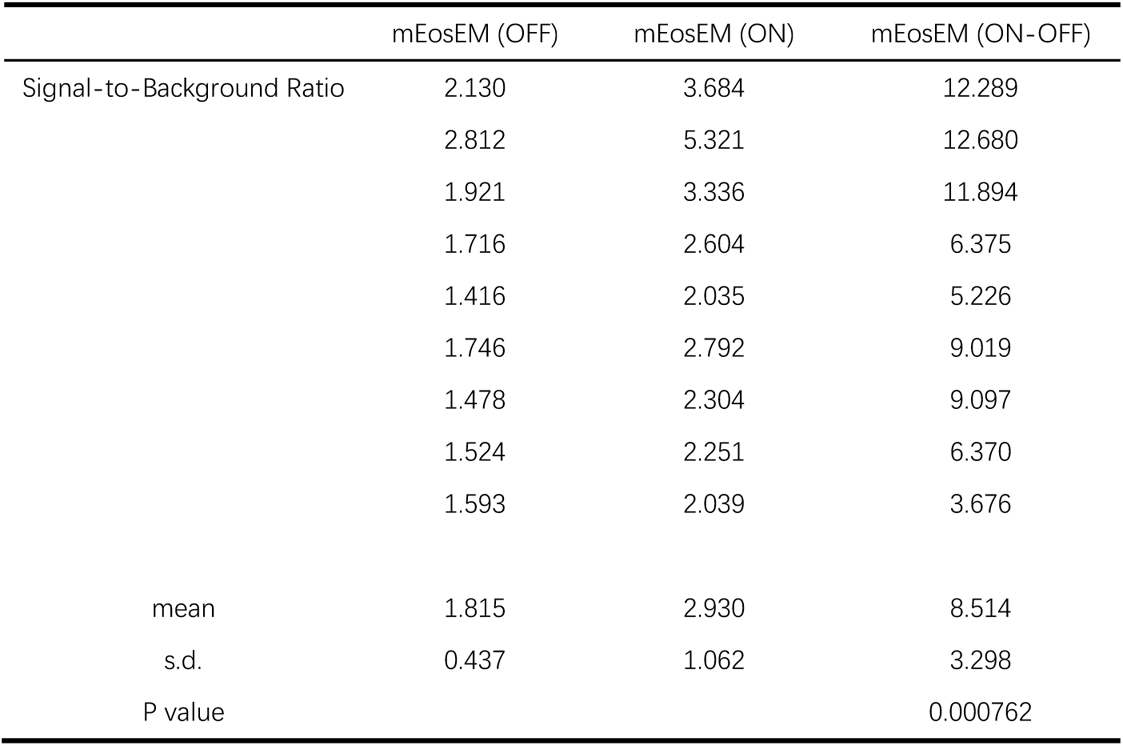
Statistics for SBR of mEosEM in OFF, ON, and ON ─ OFF images. Two-tailed t-test was performed between mEosEM (ON) and mEosEM (ON ─ OFF), n = 9.

**Table S3.** Statistics for SBR of mEosEM-E in OFF, ON, and ON ─ OFF images. Two-tailed t-test was performed between mEosEM-E (ON) and mEosEM-E (ON ─ OFF), n = 9.

|  | mEosEM-E<br>(OFF) | mEosEM-E (ON) | mEosEM-E<br>(ON-OFF) |
| --- | --- | --- | --- |
| Signal-to-Background Ratio | 2.086 | 5.785 | 28.925 |
|  | 1.651 | 4.731 | 16.408 |
|  | 1.931 | 7.015 | 34.881 |
|  | 2.020 | 7.132 | 27.124 |
|  | 1.899 | 5.406 | 38.488 |
|  | 2.978 | 8.000 | 22.849 |
|  | 2.263 | 6.488 | 28.004 |
|  | 1.966 | 6.808 | 37.050 |
|  | 2.061 | 6.655 | 31.652 |
| mean | 2.095 | 6.447 | 29.487 |
| s.d. | 0.370 | 0.990 | 7.029 |
| P value |  |  | 7.87E-06 |

**Table S4.** Statistics for SBR comparison between mEosEM and mEosEM-E in ON ─ OFF images. Two-tailed t-test was performed between mEosEM and mEosEM-E, n = 9.

|  | mEosEM | mEosEM-E |
| --- | --- | --- |
| Signal-to-Background Ratio | 12.289 | 28.925 |
|  | 12.680 | 16.408 |
|  | 11.894 | 34.881 |
|  | 6.375 | 27.124 |
|  | 5.226 | 38.488 |
|  | 9.019 | 22.849 |
|  | 9.097 | 28.004 |
|  | 6.370 | 37.050 |
|  | 3.676 | 31.652 |
| mean | 8.514 | 29.487 |
| s.d. | 3.298 | 7.029 |
| P value | 4.70E-06 |  |

**Table S5.** Statistics for fluorescence intensity comparison of RFPs after 1% OsO4 post-fixation. Two-tailed t-tests were performed between mKate2 and mScarlet, mScarlet-I, mScarlet-H, mRuby3, respectively, n = 3. Two-tailed t-tests were performed between mCherry2 and mScarlet, mScarlet-I, mScarlet-H, mRuby3, respectively, n = 3.

|  | mScarlet | mScarlet-I | mScarlet-H | mKate2 | FusionRed-MQV | mCherry2 | mRuby3 | mApple | tdTomato |
| --- | --- | --- | --- | --- | --- | --- | --- | --- | --- |
| Fluorescence |  |  |  |  |  |  |  |  |  |
| Intensity | 741.800 | 791.900 | 609.200 | 322.500 | 445.800 | 552.600 | 743.900 | 311.400 | 482.700 |
|  | 712.100 | 764.000 | 592.400 | 320.800 | 440.100 | 505.400 | 738.000 | 304.400 | 495.900 |
|  | 724.400 | 833.300 | 625.800 | 330.300 | 438.100 | 519.300 | 776.600 | 310.600 | 477.000 |
| Mean | 726.100 | 796.400 | 609.133 | 324.533 | 441.333 | 525.767 | 752.833 | 308.800 | 485.200 |
| s.d. | 14.923 | 34.868 | 16.700 | 5.066 | 3.995 | 24.255 | 20.793 | 3.831 | 9.695 |
| P value |  |  |  |  |  |  |  |  |  |
| (compared to |  |  |  |  |  |  |  |  |  |
| mKate2) | 0.000127 | 0.00150 | 0.000478 |  |  |  | 0.000423 |  |  |
| P value |  |  |  |  |  |  |  |  |  |
| (compared to |  |  |  |  |  |  |  |  |  |
| mCherry2) | 0.000714 | 0.000695 | 0.0108 |  |  |  | 0.000286 |  |  |

**Table S6.**
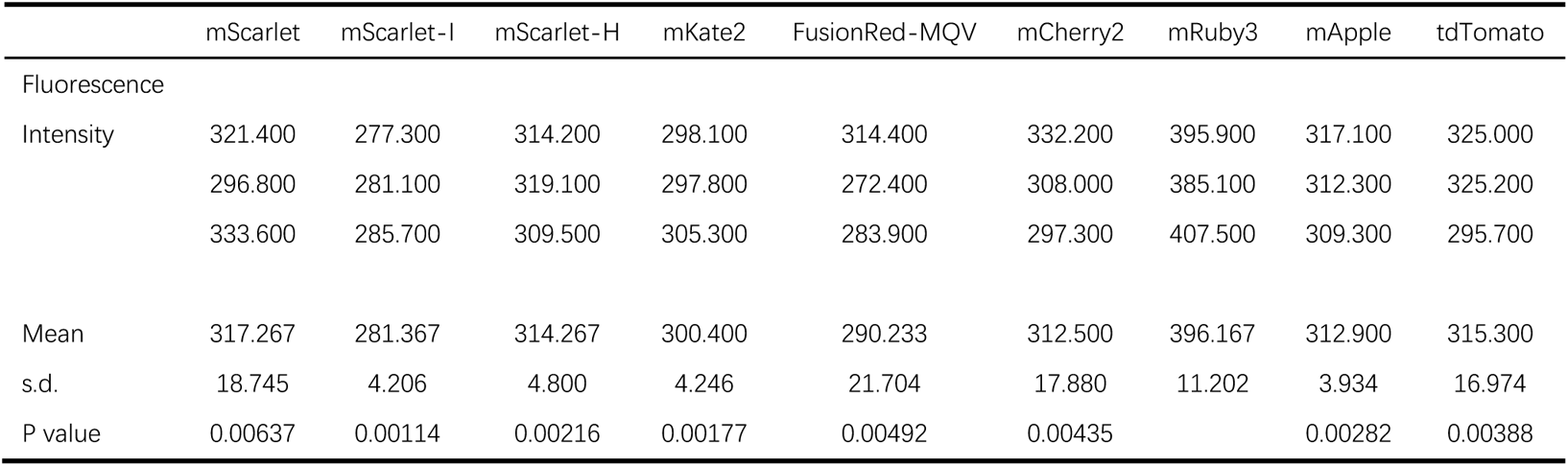
Statistics for fluorescence intensity comparison of RFPs after 1% OsO4 post-fixation and dehydration treatment. Two-tailed t-tests were performed between mRuby3 and other RFPs, n = 3.

**Table S7.** Statistics for thermostability comparison of RFPs. Two-tailed t-tests were performed between mScarlet-H and other RFPs, n = 3.

|  | mScarlet | mScarlet-l | mScarlet-H | mKate2 | FusionRed-MQV | mCherry2 | mRuby3 | mApple | tdTomato |
| --- | --- | --- | --- | --- | --- | --- | --- | --- | --- |
| Thermostability |  |  |  |  |  |  |  |  |  |
| Ratio | 0.840 | 0.682 | 0.994 | 0.071 | 0.022 | 0.214 | 0.852 | 0.550 | 0.794 |
|  | 0.842 | 0.742 | 0.959 | 0.071 | 0.021 | 0.206 | 0.856 | 0.575 | 0.728 |
|  | 0.856 | 0.750 | 0.975 | 0.066 | 0.021 | 0.205 | 0.822 | 0.603 | 0.761 |
| Mean | 0.846 | 0.725 | 0.976 | 0.069 | 0.021 | 0.208 | 0.843 | 0.576 | 0.761 |
| s.d. | 0.009 | 0.037 | 0.017 | 0.003 | 0.001 | 0.005 | 0.018 | 0.026 | 0.033 |
| P value | 0.00151 | 0.00223 |  | 8.67E-05 | 0.000109 | 5.71E-05 | 0.000840 | 7.76E-05 | 0.00199 |

**Table S8.** Statistics for fluorescence intensity comparison of RFPs after EM sample preparation. Mann-Whitney U tests were performed between mScarlet-H and each RFP, n = 106.

|  | mScarlet | mScarlet-I | mScarlet-H | mKate2 | FusionRed-MQV | mCherry2 | mRuby3 |
| --- | --- | --- | --- | --- | --- | --- | --- |
| Mean | 238.979 | 283.083 | 327.015 | 210.963 | 153.836 | 136.175 | 158.454 |
| s.d. | 135.404 | 123.250 | 178.850 | 107.039 | 58.967 | 48.306 | 77.657 |
| P value | 0.000005 | 0.144 |  | 1.78E-09 | 3.23E-21 | 1.89E-25 | 1.19E-18 |

**Table S9.** Statistics for photostability comparison of RFPs. Two-tailed t-tests were performed between mScarlet-H and other RFPs, n = 3.

|  | mScarlet | mScarlet-l | mScarlet-H | mKate2 | FusionRed-MQV | mCherry2 | mRuby3 |
| --- | --- | --- | --- | --- | --- | --- | --- |
| t1/e | 341.040 | 359.090 | 476.280 | 366.780 | 269.560 | 307.030 | 636.990 |
|  | 377.370 | 389.060 | 389.790 | 424.040 | 345.240 | 307.270 | 471.980 |
|  | 351.770 | 353.860 | 452.630 | 426.160 | 273.080 | 327.990 | 527.020 |
|  | 300.120 | 310.990 | 436.420 | 371.070 | 275.120 | 389.660 | 521.720 |
|  | 380.550 | 433.362 | 440.220 | 290.670 | 253.760 | 381.610 | 482.540 |
| Mean | 344.147 | 366.071 | 443.090 | 362.633 | 267.320 | 366.420 | 510.427 |
| s.d. | 32.610 | 45.379 | 31.644 | 55.259 | 35.600 | 40.197 | 65.429 |
| P value | 0.0470 | 0.162 |  | 0.176 | 6.37E-05 | 0.0508 | 0.0299 |

**Table S10.**
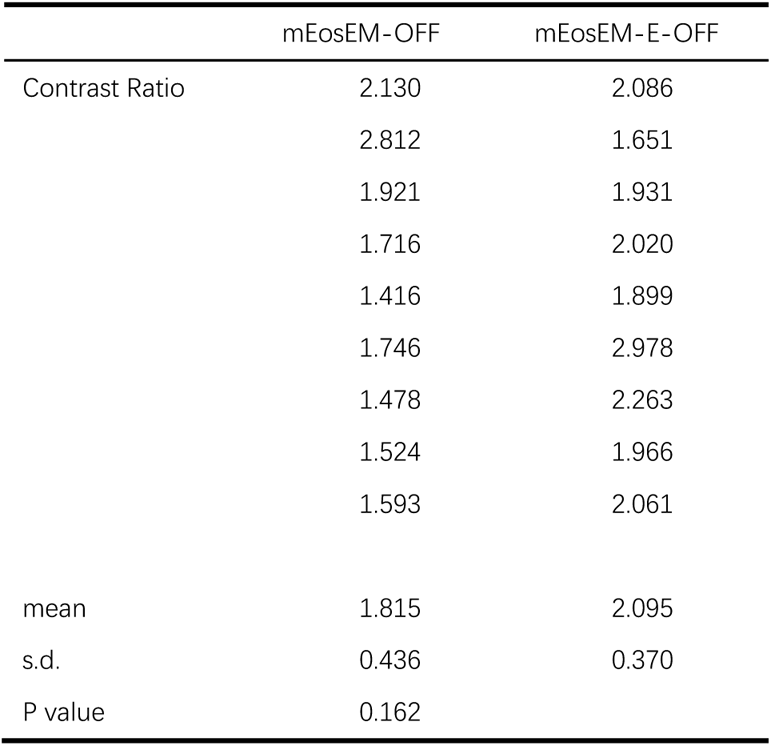
Statistics for SBR comparison between mEosEM and mEosEM-E in OFF images. Two-tailed t-test was performed between mEosEM and mEosEM-E, n = 9.

**Table S11.**
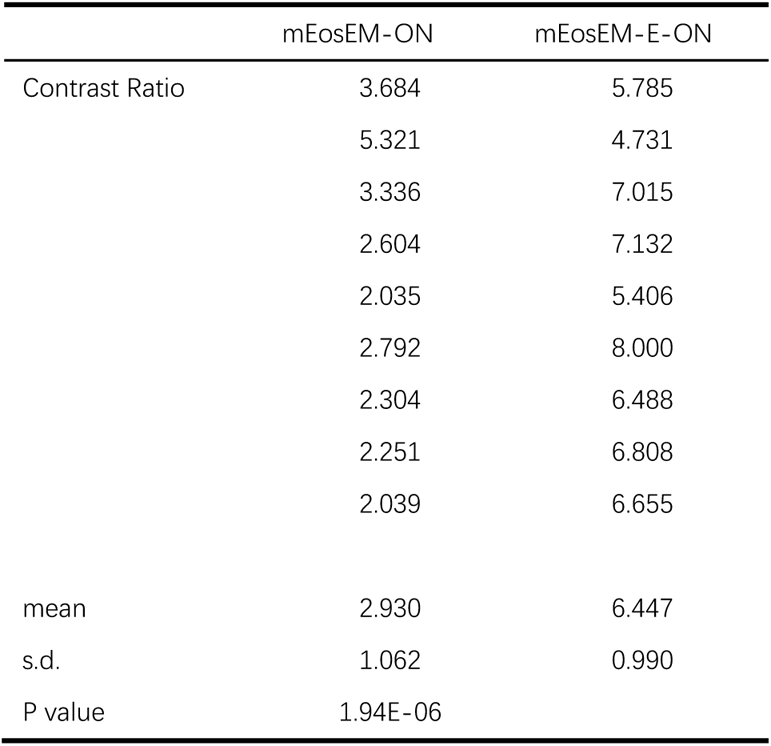
Statistics for SBR comparison between mEosEM and mEosEM-E in ON images. Two-tailed t-test was performed between mEosEM and mEosEM-E, n = 9.

|  | mEosEM-ON | mEosEM-E-ON |
| --- | --- | --- |
| Contrast Ratio | 3.684 | 5.785 |
|  | 5.321 | 4.731 |
|  | 3.336 | 7.015 |
|  | 2.604 | 7.132 |
|  | 2.035 | 5.406 |
|  | 2.792 | 8.000 |
|  | 2.304 | 6.488 |
|  | 2.251 | 6.808 |
|  | 2.039 | 6.655 |
| mean | 2.930 | 6.447 |
| s.d. | 1.062 | 0.990 |
| P value | 1.94E-06 |  |

**Table S12.** Statistics for fluorescence intensity comparison of RFPs in live cells. n = 3.

|  | mScarlet | mScarlet-l | mScarlet-H | mKate2 | FusionRed-MQV | mCherry2 | mRuby3 | mApple | tdTomato |
| --- | --- | --- | --- | --- | --- | --- | --- | --- | --- |
| Fluorescence |  |  |  |  |  |  |  |  |  |
| Intensity | 2928 | 3256 | 1453 | 1406 | 1568 | 1752 | 3011 | 3364 | 4674 |
|  | 2906 | 3269 | 1457 | 1344 | 1763 | 1715 | 3100 | 3423 | 4637 |
|  | 2887 | 3449 | 1348 | 1401 | 1719 | 1781 | 3104 | 3465 | 4643 |
| Mean | 2907 | 3324.667 | 1419.333 | 1383.667 | 1683.333 | 1749.333 | 3071.667 | 3417.333 | 4651.333 |
| s.d. | 20.51828 | 107.8718 | 61.80885 | 34.44319 | 102.2758 | 33.08071 | 52.57693 | 50.73789 | 19.85783 |

**Table S13.** Statistics for fluorescence intensity comparison of RFPs after pre-fixation. n = 3.

|  | mScarlet | mScarlet-l | mScarlet-H | mKate2 | FusionRed-MQV | mCherry2 | mRuby3 | mApple | tdTomato |
| --- | --- | --- | --- | --- | --- | --- | --- | --- | --- |
| Fluorescence |  |  |  |  |  |  |  |  |  |
| Intensity | 1490 | 1819 | 955.6 | 651.3 | 1117 | 974.2 | 1995 | 703.9 | 1860 |
|  | 1483 | 1903 | 966.4 | 639.8 | 1228 | 1019 | 2198 | 715.2 | 2041 |
|  | 1450 | 1841 | 924.8 | 642.8 | 1178 | 1041 | 2000 | 803.6 | 1745 |
| Mean | 1474.333 | 1854.333 | 948.9333 | 644.6333 | 1174.333 | 1011.4 | 2064.333 | 740.9 | 1882 |
| s.d. | 21.36196 | 43.55839 | 21.58642 | 5.965177 | 55.59077 | 34.04233 | 115.7857 | 54.59295 | 149.2213 |

**Table S14.** Statistics for fluorescence intensity comparison of RFPs after dehydration treatment. Two-tailed t-tests were performed between mRuby3 and other RFPs, n = 3.

|  | mScarlet | mScarlet-l | mScarlet-H | mKate2 | FusionRed-MQV | mCherry2 | mRuby3 | mApple | tdTomato |
| --- | --- | --- | --- | --- | --- | --- | --- | --- | --- |
| Fluorescence |  |  |  |  |  |  |  |  |  |
| Intensity | 429.100 | 583.500 | 648.900 | 685.600 | 518.800 | 938.800 | 1553.000 | 1153.000 | 680.100 |
|  | 442.600 | 491.500 | 604.900 | 681.800 | 487.400 | 878.500 | 1615.000 | 917.600 | 650.000 |
|  | 440.500 | 780.200 | 639.500 | 665.700 | 483.300 | 823.900 | 1525.000 | 865.900 | 549.900 |
| Mean | 437.400 | 618.400 | 631.100 | 677.700 | 496.500 | 880.400 | 1564.333 | 978.833 | 626.667 |
| s.d. | 7.264 | 147.480 | 23.172 | 10.565 | 19.421 | 57.474 | 46.058 | 153.032 | 68.164 |
| P value | 0.000420 | 0.00461 | 8.06E-05 | 0.000526 | 0.000102 | 0.000119 |  | 0.0157 | 0.000101 |

**Table S15.** Statistics for SBR comparison of RFPs after EM sample preparation. Mann-Whitney U tests were performed between mScarlet-H and each RFP, n = 106.

|  | mScarlet | mScarlet-I | mScarlet-H | mKate2 | FusionRed-MQV | mCherry2 | mRuby3 |
| --- | --- | --- | --- | --- | --- | --- | --- |
| Mean | 1.791 | 1.910 | 2.034 | 1.706 | 1.524 | 1.456 | 1.541 |
| s.d. | 0.385 | 0.335 | 0.495 | 0.307 | 0.181 | 0.147 | 0.237 |
| P value | 0.000009 | 0.182 |  | 6.14E-09 | 8.95E-22 | 6.34E-27 | 1.64E-18 |

